# A universal polyphosphate kinase powers *in vitro* transcription

**DOI:** 10.1101/2025.03.21.644484

**Authors:** Ryusei Matsumoto, Takayoshi Watanabe, Eishin Yamazaki, Ako Kagawa, Liam M. Longo, Tomoaki Matsuura

**Author notes:** The authors contributed equally to this study (Ryusei Matsumoto, Takayoshi Watanabe).

## Abstract

Polyphosphate kinases (PPKs) catalyze phosphoryl transfer between polyphosphates and nucleotides. Polyphosphates are a cost-effective source of phosphorylating power, making polyphosphate kinases an attractive enzyme for nucleotide production and regeneration. However, at present, applications that require the simultaneous utilization of diverse nucleotides, such as RNA transcription, remain out of reach due to the restricted substrate profiles of known PPK enzymes. Here, we present the discovery and characterization of a universal PPK capable of efficiently phosphorylating all eight common ribonucleotides: purines and pyrimidines, monophosphates and diphosphates to triphosphates. Under optimal reaction conditions, approximately 70% triphosphate conversion was observed for all common ribonucleotides, with only limited over-phosphorylation. At elevated temperatures, however, production of adenine-capped polyphosphates up to a 30-mer was achieved. An analysis of mutant and chimeric enzymes revealed a rugged functional landscape, particularly for non-adenine nucleotides. Finally, we demonstrated the biotechnological potential of a universal PPK enzyme with a one-pot assay for PPK-powered *in vitro* transcription.

## Introduction

In 1956, Kornberg reported the discovery of polyphosphate kinases, enzymes that transfer phosphate groups between nucleotides and polyphosphate (PolyP) ^1,2^. Inspired by the structural simplicity of PolyP, its facile production from orthophosphate by heating, and its role in intracellular phase phenomena ^3^, Kornberg hypothesized that PolyP served as a primitive source of phosphorylating power ^4,5^. Nearly 50 years later, Kornberg reported a second family of polyphosphate kinases (PPK2) ^6,7^ belonging to the ancient and ubiquitous P-Loop NTPase enzyme lineage ^8,9^. Whereas PPK1 favors PolyP elongation, PPK2 favors nucleotide phosphorylation and is well suited for energy regeneration, in which a pool of nucleotide triphosphates is maintained to drive catalysis ^4,10^. The first PPK2 enzymes to be characterized could phosphorylate either nucleotide monophosphates (Class II) or diphosphates (Class I). However, in 2014, the discovery of Class III PPK2 enzymes ^11^ that can convert AMP to ATP further expanded the scope of energy regeneration by PolyP ^6,7^, which is cheaper than more popular stores of phosphorylating power such as pyruvate or creatine phosphate ^12^. Increasingly, PPK2 is emerging as the key enzyme to developing a robust and cost-effective PolyP-driven energy regeneration system with broad biotechnological applicability, including *in vitro* chemical production^13^.

While metabolic and genome engineering of microbes for chemical production has advanced rapidly, the unpredictable, difficult-to-control nature of living organisms can limit the utility of these approaches^14^. Consequently, various *in vitro* platforms for chemical production have been developed – including reactions of immobilized enzymes, cascades mediated by multiple purified enzymes, and reactions carried out in crude cell extracts^15^, many of which require energy regeneration. Proof-of-principle demonstrations of PPK2-powered *in vitro* chemical synthesis indicate significant progress towards realizing the goal of PolyP-driven regeneration ^6,7 16–18^, particularly for simple reactions that require a *single* nucleotide. Excitingly, a broad-specificity PPK2 enzyme that efficiently phosphorylates both adenine and guanine ribonucleotides (but not pyrimidines) was recently identified ^19^ and applied to *in vitro* translation^20^, which requires *two* nucleotide regeneration, ATP and GTP. However, reaction systems that require all *four* common nucleotides (ATP, GTP, CTP, and UTP) still remain out of reach. While ATP and GTP often serve as an energy source, UTP and CTP are also known to play important roles in metabolism. For example, UTP and CTP can activate sugars^21^ and lipid precursors^22^, respectively, that precede the production of various chemicals. Thus, a single enzyme capable of regenerating all common ribonucleotides would facilitate the design of on-demand *in vitro* metabolic pathways that bridge diverse nucleotide requirements. Does such a universal PPK2 (uPPK2) enzyme that can efficiently phosphorylate all four common ribonucleotides exist?

Here, we report the discovery of a uPPK2 and demonstrate its utility by powering *in vitro* transcription. Four Class III PPK2 enzymes were selected and tested for their universal nucleotide phosphorylation activity. Despite close evolutionary relationships, only one enzyme – the PPK2 from *Mangrovibacterium marinum*, designated MAN – was able to phosphorylate all four common nucleotide monophosphates (AMP, GMP, CMP, UMP) and diphosphates (ADP, GDP, CDP, UDP) to their corresponding triphosphate (ATP, GTP, CTP, UTP), achieving ∼70% conversion in each case. At elevated temperatures, however, adenine polyphosphates up to AP30, the longest nucleotide polyphosphate reported to date ^23–25^, could be produced. Key residues involved in the universal activity of MAN were investigated, in which all analyzed mutants lost universal activity, revealing a rugged functional landscape. Finally, we demonstrate that MAN-mediated, one-pot synthesis of NTPs from NMPs can directly power *in vitro* transcription (IVT). We find that enzyme promiscuity can be a design principle for constructing simple reaction systems – a notable parallel to the proposed role of promiscuous enzymes in primitive biology^26,27^. Given the simplicity of MAN-based nucleotide phosphorylation, we envision significant potential for MAN and similar enzymes in supporting diverse *in vitro* metabolic pathways ^15^.

## Results

### Sequence selection strategy

An ideal uPPK2 should phosphorylate both pyrimidines and purines, and produce triphosphates from both monophosphate and diphosphate nucleotides – eight substrates in total. To find a naturally occurring uPPK2, we focused on Class III PPK2 enzymes^10,11,19^. Although an analysis of PPK2 gene complement across prokaryotes challenges a class-based heuristic for functional prediction ^28,29^, Class III enzymes have two attractive properties: First, they have been demonstrated to convert NMPs to NTPs either via two successive phosphoryl transfer reactions or by pyrophosphoryl transfer ^10,11,16,19,24^. Second, some Class III enzymes can accept both adenine and guanine nucleotides due to a key substrate-broadening mutation, an Asn at position 138 (numbering based on the PPK2 enzyme from *Cytophaga hutchinsonii*, referred to here as CHU^19^). By flipping the side-chain carboxamide of Asn, the enzyme can accommodate both the C6 amino group of adenine and the C6 carbonyl group of guanine (**Figure 1A**). We hypothesized that a uPPK2 enzyme may employ the same mechanism to achieve broad purine specificity. And, with its ability to act as either a hydrogen bond donor or acceptor, N138 may also support the binding of both pyrimidines in the right context. For these reasons, we focused our search on Class III PPK2 enzymes with an Asn at position 138.

A gene tree of Class III PPK2 reveals that enzymes with N138 belong to a limited number of clades (**Figure 1B**). The major clade, to which most N138 enzymes belong, also contains the previously characterized CHU enzyme ^19^. In addition to the above, the selection of PPK2 genes for functional profiling followed three simple criteria: First, PPK2 sequences should be of canonical length, roughly 290 residues ^19,28^. Second, genes should come from mesophiles for compatibility with various enzyme assay conditions, including cell-free protein synthesis^16,30,31^. Finally, while around 30% of bacterial species are known to have multiple copies of PPK2 genes ^28^, only species harboring one PPK2 gene were considered, to exclude cases where a division of labor between two or more enzymes may occur^3,6^. In total, four genes were selected from the major N138 clade of Class III (**Figure 1B** and **C, Table S2**): The PPK2 enzymes from *Pedobactor sp*. (PED), *Mangrovibacterium marinum* (MAN), *Arenitalea lutea* (ARE), and *Leeuwenhoekiella palythoae* (LEE). The pairwise amino acid identities of the selected enzymes range from 44.9-68.0%.

**Figure 1.**
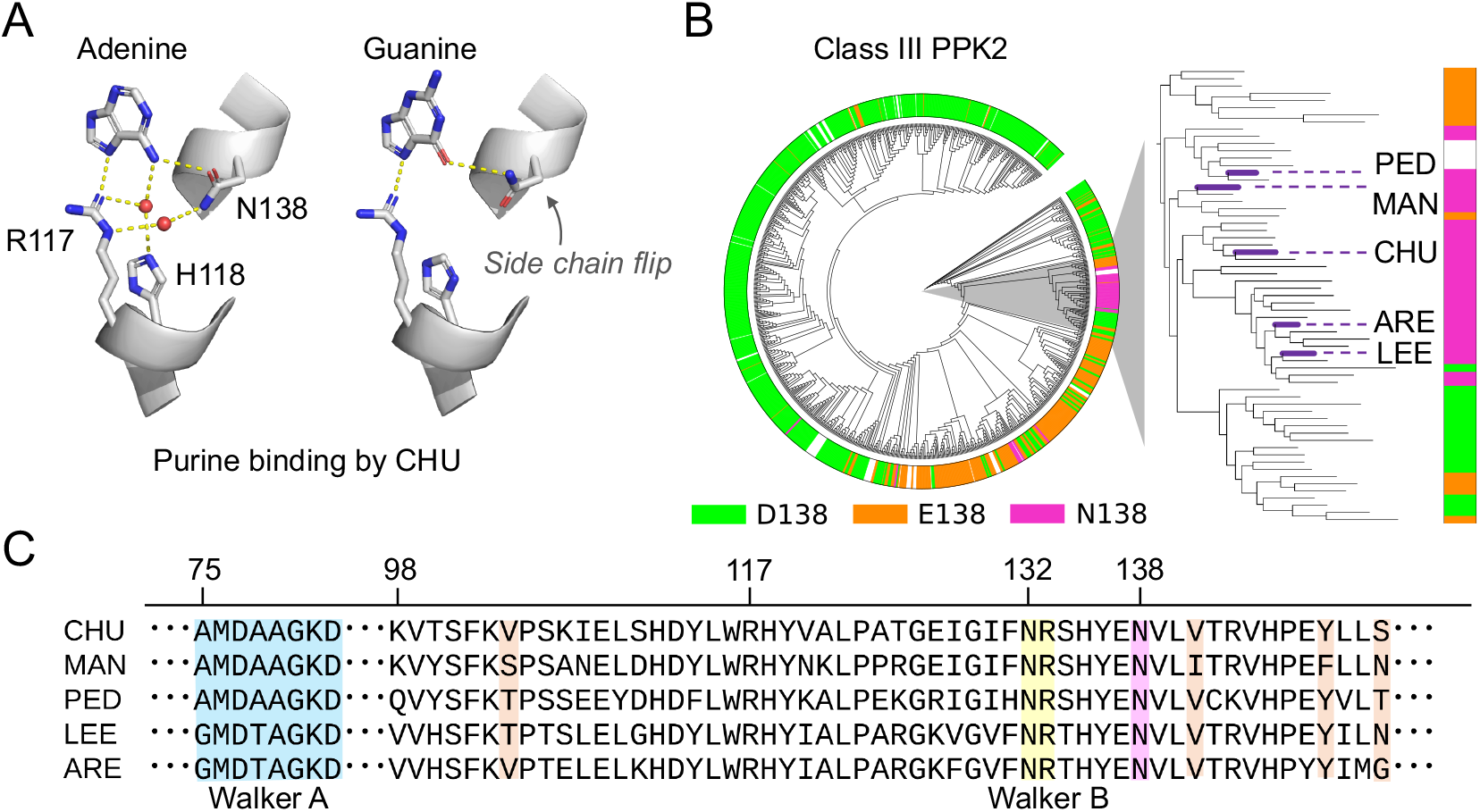
Gene selection strategy. **A**. Side-chain flipping of N138 to mediate binding of adenine (PDB ID 6AN9) and guanine (PDB ID 6ANH)^19^. The primary polar interactions with adenine involve R117, H118, and N138, which form a network of hydrogen bonds with each other, the base, and two water molecules^19,32^. The hydrogen bonding network upon binding guanine is unclear, as the lower resolution of the crystal structure complicates the interpretation of absent waters. **B**. Class III PPK2 gene tree (left) and a zoom-in on the major N138 clade (right). Genes selected for experimental characterization, as well as the previously characterized CHU enzyme, are indicated. **C**. Multiple sequence alignment (CHU numbering) of selected constructs highlighting key functional regions^19,33–35^, including the Walker A (blue), Walker B (yellow), and N138 (magenta). All selected sequences contain a variation of the Walker A motif, canonically GxxxxGK[TS], that lacks a conserved Ser/Thr at the last position, as is common among PPK2 enzymes^8^. Residues of the base binding pocket that differ between CHU and MAN are highlighted in orange (see also **Figure 4**).

### MAN is a universal PPK2

Phosphoryl transfer activity of each selected PPK2 was quantified by HPLC (**Figure 2A**) for eight nucleotides (four mono- and four di-phosphate nucleotides) and three PolyP chain lengths (10, 60 and 700 phosphate residues on average). Although the resulting functional profiles, reported here as fractional conversion to NTP, are relatively idiosyncratic (**Figure 2B**), several general trends emerge: First, all enzymes tested are able to catalyze phosphoryl transfer to both mono- and diphosphates, consistent with other Class III PPK2 enzymes. Second, PolyP-60 is preferred by all four enzymes, though the influence of PolyP length is not uniform and even adenine nucleotide utilization, which is an enduring feature of the PPK2 family, is sensitive to PolyP length (e.g., MAN in Poly-60 vs. PolyP-700). Finally, purines are generally preferred over pyrimidines. However, despite all proteins having N138, GTP production varied greatly, due in part to over-phosphorylation (**Figure S3**). ARE and LEE are the most closely related enzymes (68.0% identity), yet PED and ARE (45.6% identity) exhibited the most similar functional profiles – greater sequence similarity does not necessarily imply greater similarity in functional profiles, at least at these identity ranges.

**Figure 2.**
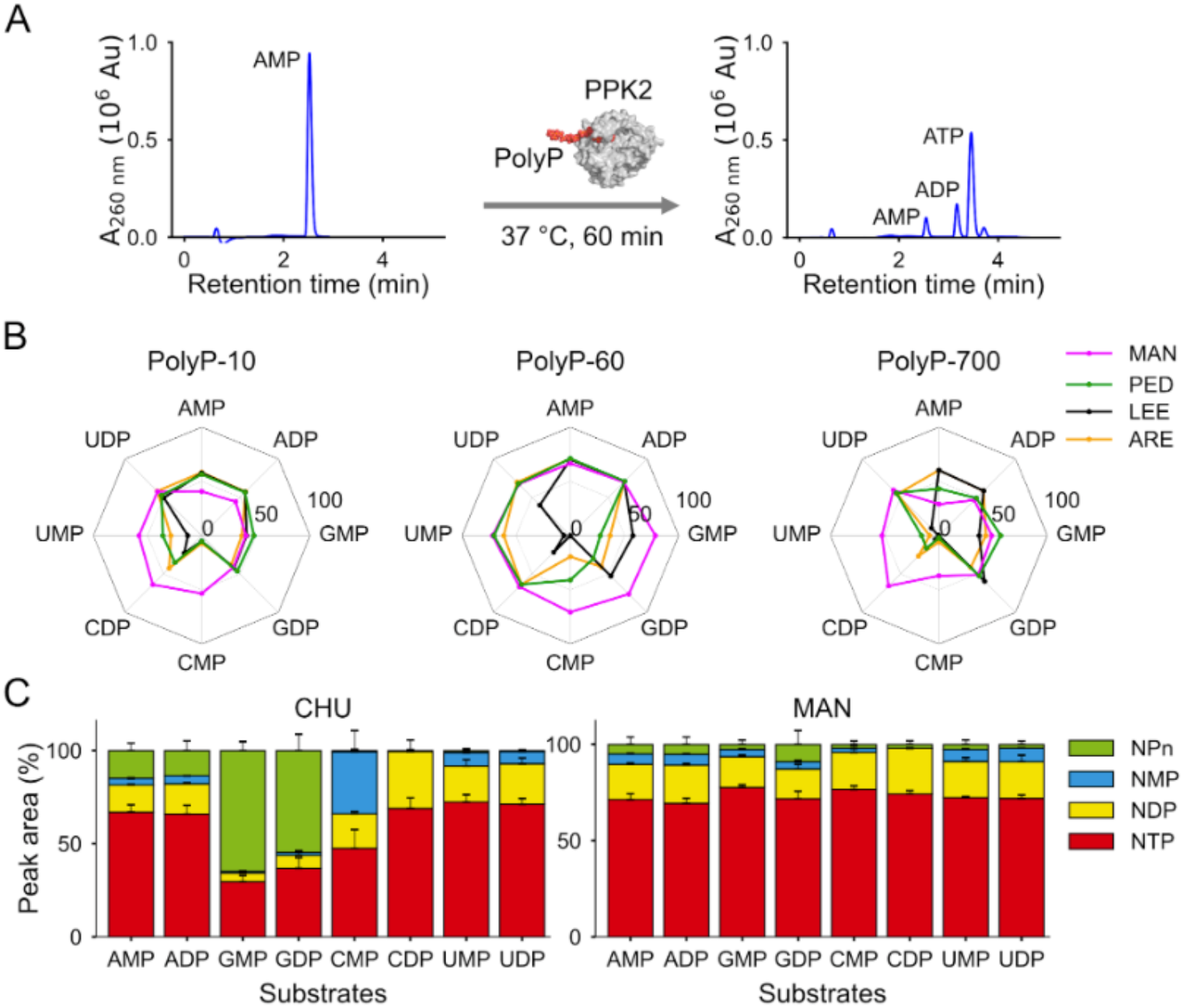
Substrate profiling of selected Class III PPK2 enzymes. **A**. Overview of the enzyme profiling assay. The product distribution was analyzed by HPLC after 60 min incubation at 37 °C. Refer to **Figure S2** for additional details about the HPLC analysis. **B**. Percent conversion to NTP for selected enzymes (18.9 ^μ^M) reacting with the indicated nucleotide (4 mM) and PolyP chains of varying length (65 mM phosphate units) in the presence of 10 mM Mn^2+^. Refer to **Figure S3** for product distributions. **C**. Product distribution of MAN and CHU under the same conditions as in panel B. Plotted values are the average of 4 independent runs. Error bars are standard deviations.

While most enzymes failed to phosphorylate one or several nucleotides, only MAN catalyzed significant triphosphate production for all eight nucleotides and all three PolyP lengths (**Figure 2B**). In reactions with PolyP-60, where MAN is the most efficient, 67-79% NTP conversion for NMPs, NDPs, purines, and pyrimidines was achieved. Moreover, an analysis of MAN-catalyzed NTP production with PolyP-60 (**Figure 2C**) reveals a remarkably consistent product distribution for all nucleotides, with only minor production of over-phosphorylated nucleotides (NPn). Recently, CHU has emerged as a valuable resource for biotechnology applications, highlighting the potential utility of broad-specificity PPK2 enzymes for nucleotide regeneration ^18,20,36^. Noted for having a relatively high *k*cat/K_M_ of 10^3^ s^-1^M^-1^ for GMP and GDP, CHU is an ideal point of comparison for MAN. We observe that under the same assay conditions, CHU produces significant amounts of GPn (guanine nucleotides with four or more phosphate groups) and has weak activity against CMP. These limitations are similar to those of the other enzymes tested: PED and ARE exhibit weak activity against CMP, GDP, and GMP, while LEE exhibits no activity against CMP and UMP, with at best weak activity against CDP and UDP. Based on these data, we designate MAN a uPPK2 enzyme – unique among the other proteins characterized from the major N138 clade of PPK2 Class III enzymes.

### Controllable production of ATP and APn

In addition to PolyP length, MAN activity may also be modulated by temperature^11,37,38^, dication availability^11,19,37,39^, and PolyP concentration^37,38,40^ (**Figure 3**). Optimal ATP synthesis was observed between 37-45 °C using AMP and PolyP-60 as substrates (**Figure 3A**, left). At lower temperatures, the rate of the reaction is reduced, and the proportion of unreacted AMP increases. At 55 °C, however, the HPLC chromatogram is characterized by a broad, wavy tail at high retention times (**Figure 3A**, right). Previously, this wavy pattern was identified as over-phosphorylated nucleotides by LC-MS/MS ^23–25^, indicating that the product distribution has shifted to over-phosphorylation. Careful inspection of the chromatogram reveals nucleotides with 30 or more attached phosphate groups. Over-phosphorylation at higher temperatures may be due to partial unfolding and/or increased flexibility of the enzyme^41^ and this interpretation is consistent with *M. marinum* (the parent organism of MAN) having an optimal growth temperature of 33 °C^42^. To our knowledge, AP30 is the longest reported adenine nucleotide to date, with AP9 holding the previous record ^23^. The biological occurrence of adenine-capped PolyP molecules (such as AP10 and AP30) remains to be elucidated^23,25^. Compared to AMP, only modest amounts of GPn formation occur at 55 °C, and over-phosphorylation of pyrimidines is not observed (**Figure S4**).

**Figure 3.**
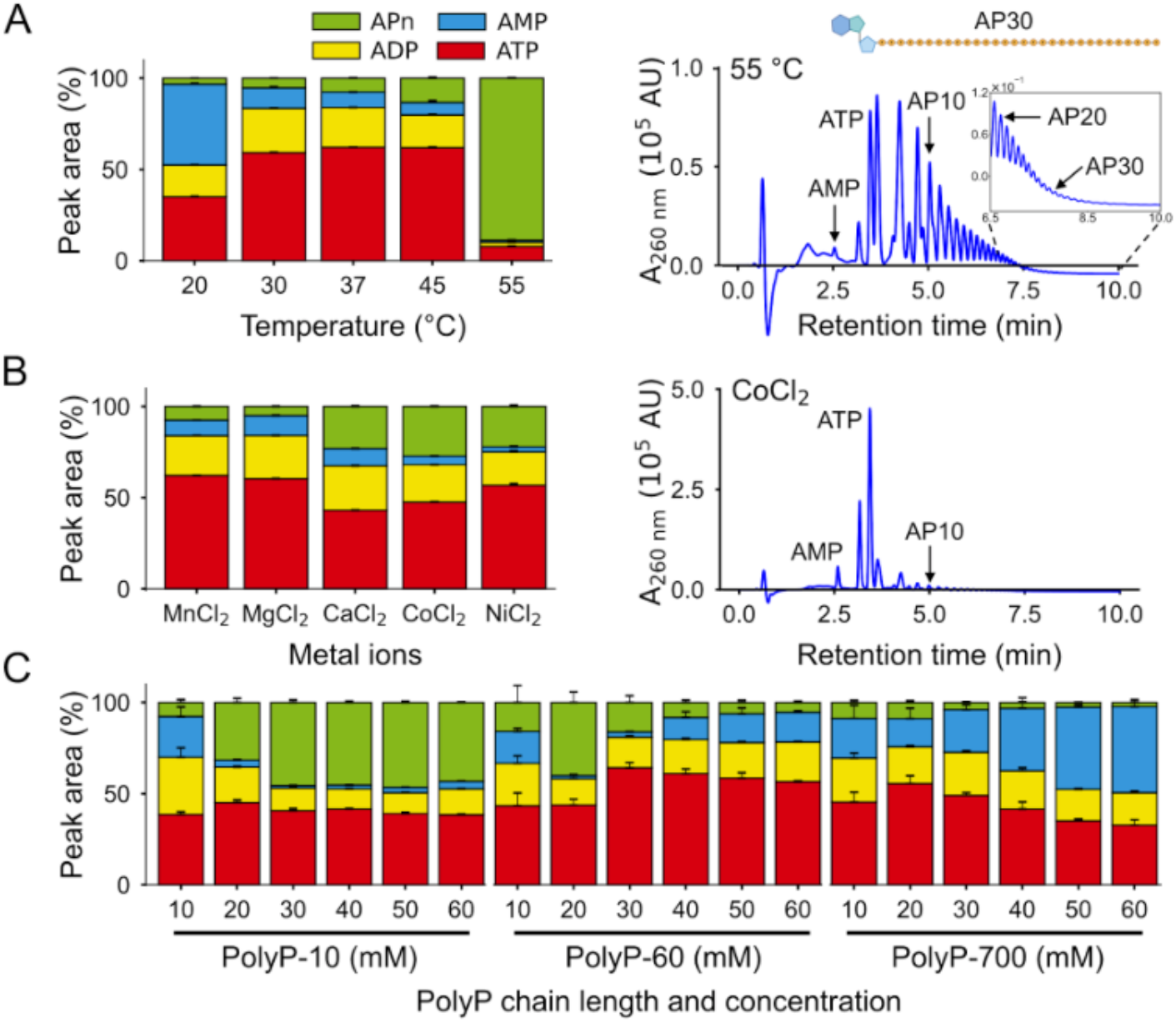
Effect of temperature (**A**), metal ion (**B**), and PolyP concentration (**C**) on MAN catalysis. Unless otherwise indicated, reactions were performed at 37 °C and contained 4 mM AMP, 65 mM phosphate units of PolyP-60, and 10 mM Mn^2+^. Plotted values are the average of 4 independent runs. Error bars are standard deviations.

MAN, like other PPK2 enzymes^19,39^, can accept diverse divalent metal ions (**Figure 3B**). The highest NTP conversions were observed with Mn^2+^ and Mg^2+^, which have roughly equivalent product distributions. Conversely, Ca^2+^, Co^2+^, and Ni^2+^ promote APn production, though not as much as increased temperature. PolyP concentration effects are also significant and PolyP-length dependent: Whereas high concentrations of PolyP-700 and PolyP-60 suppress reactivity, decreasing the concentration of PolyP-10 suppresses over-phosphorylation. A ladder of decreasing APn sizes is observed, suggesting that APn is produced by sequential reactions and not direct attachment of the nucleotide and PolyP molecules. In short, the extent of over-phosphorylation by MAN can be controllably suppressed or exploited by assay conditions.

### MAN kinetic characterization

Kinetic parameters were determined for MAN (**Table 1, Figure S5**) in the presence of Mg^2+^ at 30 °C. The dication Mg^2+^ was selected because it is a common component of *in vitro* transcription and translation systems. 30 °C was chosen to facilitate direct comparison with prior work on other Class III PPK2 enzymes ^19^. Binding of diphosphates is preferred over monophosphates, except for guanine nucleotides, where the trend is reversed. Turnover numbers, on the other hand, are consistently higher for monophosphates than for diphosphates, with an 8% (guanine) to 800% (uracil) increase. Binding of purine monophosphates (K_M_ = 4.6-7.7 mM) is preferred over pyrimidine monophosphate (K_M_ = 26.2-47 mM) and turnover numbers show a clear preference for purines (*k*_cat_ = 29.7-74.6 s^-1^) over pyrimidines (*k*_cat_ = 0.36-8.9 s^-1^) as well. In total, second-order rate constants (*k*_cat_/K_M_) are higher for purines than pyrimidines, and all nucleotides except CMP and CDP have a second-order rate constant greater than 10^2^ s^-1^M^-1^. AMP has the highest catalytic efficiency of 1.6 × 10^4^ s^-1^M^-1^, likely reflecting the primary biochemical role of this enzyme. Catalytic efficiencies of 10^3^-10^4^ s^-1^M^-1^ for purine nucleotides are consistent with previous studies of Class III PPK2 enzymes, which range from 1.0 × 10^2^ - 2.1 × 10^5^ s^-1^M^-1 19^. To our knowledge, a full characterization of pyrimidine catalytic efficiencies has not been reported. The kinetic characterization confirms activity against all substrates while also revealing clear nucleotide preferences.

**Table 1.**
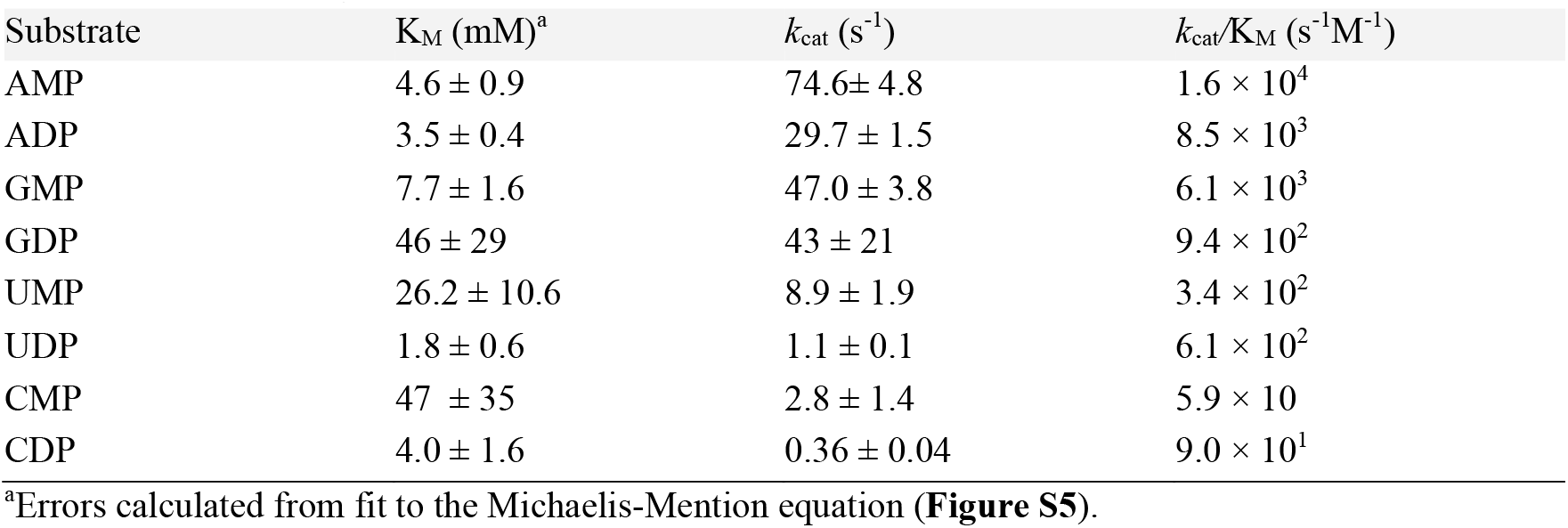
MAN kinetic parameters.

### N138 contributes to utilization of all nucleotides

N138 is thought to facilitate binding and catalysis of both adenine and guanine nucleotides (**Figure 1A**). A mutational analysis of N138 in MAN (**Figure 4A** and **B**), however, reveals a more complex picture: At high enzyme concentrations and long incubation times (10 ^μ^M and 60 min; **Figure S6A** and **B**) mutation to Ala, Asp, or Glu resulted in only modest differences in NTP production, with N138 being the most active overall. To better assess differences in enzymatic rates, the enzyme concentration was reduced to 6 ^μ^M and incubation times were reduced to either 10 min (CMP and CDP) or 5 min (all other nucleotides) (**Figure 4B, Figure S6C**). Under these conditions, mutation of N138 resulted in a non-uniform decrease in NTP production for all nucleotides tested, with AMP being the most robust. Mutation to Ala and Glu preferentially retained GMP and CDP activity, respectively, while mutation to Asp was generally the most disruptive. In all cases, decreased NTP production was due to reduced activity and not over-phosphorylation. Given the proximity of N138 to the base (**Figure 1A, Figure 4A**) and the non-uniform loss of activity across nucleotides (**Figure 4B**), we conclude that this position contributes to the broad substrate profile of MAN, as initially hypothesized^19^.

**Figure 4.**
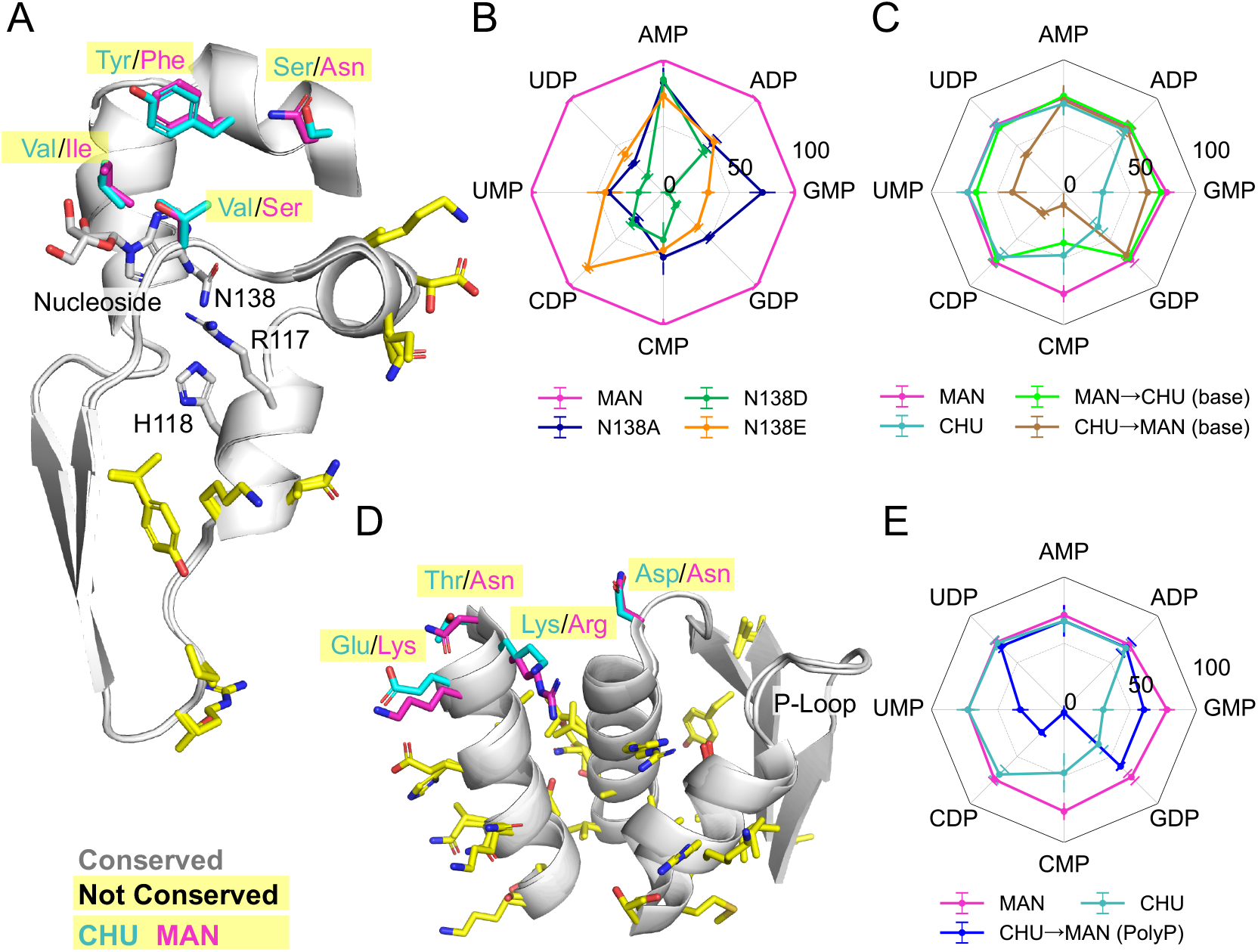
Mutation analysis of MAN and CHU. **A**. Overlay of the base binding pockets of MAN (AlphaFold3 structure AF-A0A2T5C445-F1-v4)^44^ and CHU (PDB ID 6AN9)^19^. Conserved residues other than H118, R117, and N138 that mediate binding are not drawn. Sequence differences in the primary base binding pocket are drawn as cyan and magenta sticks for CHU and MAN, respectively. Sequence differences that point away from the base binding pocket and were not characterized further are shown as yellow sticks. Additional sequence information can be found in **Figure 1C. B**. Characterization of position 138. Note that activities are normalized to MAN activity, indicated by the fuchsia outer ring. **C**. Characterization of base binding site chimeras. Refer to the main text for more details. **D**. Overlay of the polyphosphate binding track, following the coloring conventions in panel A. **E**. Characterization of polyphosphate binding site chimera. Panels C and E report the percentage of NTP production. Plotted values are the average of 4 independent runs. Error bars are standard deviations.

### Truncation of the C-terminal α-helix promotes purine utilization and suppresses pyrimidine utilization

The active site of PPK2 enzymes is centered on the Walker A motif, which adopts the canonical P-loop conformation^19,33–35^. As with thymidylate kinases^43^, an α-helix hairpin “hood” rests atop the P-loop. Upon oligomerization, the C-terminal α-helix of one protomer interacts with the hood of an adjacent protomer (**Figure S7A**), potentially influencing the catalytic properties of the enzyme. Previously, truncating the C-terminal α-helix of CHU resulted in improved purine utilization ^19^, a result reproduced here (**Figure S7B**). Truncation of the C-terminal α-helix in LEE and PED produced similar results (**Figure S7B**). The MAN truncation variant failed to express and could not be characterized. These data suggest that truncation is a general strategy for designing purine-preferring Class III PPK2 enzymes (in cases where the resulting protein is well behaved) but at the expense of pyrimidine utilization. Class III PPK2 enzymes that naturally lack the C-terminal α-helix (*e*.*g*., A0A559PSS6), although uncommon, may represent the natural exploration of this engineering approach. The exact mechanism of truncation is unclear but may relate to changes in the oligomerization state affecting conformational dynamics. Analytical ultracentrifugation of full-length and truncated variants of CHU, LEE, and PED (**Figure S8**) demonstrates a transition from a dimer to a monomer upon truncation (**Table S1**).

### Base binding preferences are encoded in the base binding pocket and beyond

What residues underlie the broad specificity of PPK2 enzymes, and can we explain the functional differences between MAN and CHU? In the region of the base binding pocket, just four residues (positions 104, 141, 148, and 151; CHU numbering) differ between MAN and CHU (**Figure 4A**, cyan and magenta sticks; **Figure 1C**, orange highlight). Although these residues do not make direct interactions with the base in the most common adenine binding mode (represented in **Figure 4**), an alternative binding mode found in Class I PPK2 (**Figure S9A** and **B**) has the base flipped up^34^, potentially within interaction distance of these residues. Other differences between MAN and CHU (**Figure 4A**, yellow sticks) are more distant and have their side chains oriented away from the base binding pocket. To better understand the determinants of substrate specificity, these four positions in the MAN binding site were mutated to the corresponding CHU residue and vice versa. MAN→CHU (base), where the parent enzyme is MAN, shows a contraction of the substrate profile, with activity against CMP becoming CHU-like and activity against guanine nucleotides largely unchanged. On the other hand, CHU→MAN (base) (**Figure 4C, Figure S9C**), where the parent enzyme is CHU, shows improved activity against GMP and GDP but reduced activity against pyrimidines (**Figure 4C, Figure S9C**). Neither of the chimeric constructs resulted in a complete conversion of base binding preferences.

The partial conversion of MAN and CHU activities by binding site chimeras suggests that base binding preferences are determined in part by non-local interactions, for example, through long-range dynamics (consistent with the truncation analysis) or the positioning of the polyphosphate substrate. To test the latter possibility, residues along the path of the polyphosphate chain (as inferred from PDB ID 5LLF; **Figure 4D**) in CHU were mutated to be MAN-like. The resulting chimera, CHU→MAN (PolyP), was more active against GMP and GDP but lost activity against the pyrimidines UMP, CDP, and CMP (**Figure 4E, Figure S9C**). The MAN→CHU (PolyP) construct was degraded in *E. coli* and could not be purified. Taken together, these data suggest that the substrate preferences of Class III PPK2 enzymes – particularly for nucleotides other than ATP – are encoded by residues distributed across the protein, and not simply near the base binding pocket.

### uPPK2-powered in vitro transcription

Previously, PPK2 was used for nucleotide recycling^16,20,30,31^; specifically, for the regeneration of ATP and GTP from ADP/AMP and GDP, respectively. With the ability to phosphorylate all four ribonucleotides, MAN represents a significant new opportunity for the biotechnological application of PPK2 enzymes. To underscore this point, we established an *in vitro* transcription (IVT) reaction system powered by MAN-derived NTPs. Our goal was to minimize costs, so even though PolyP-10 has a lower NTP conversion efficiency than PolyP-60, it was selected for the IVT assay due to its lower price. Given the differences in catalytic efficiency between purine monophosphates and pyrimidine monophosphates (**Table 1**), two reaction schemes were considered: A one-step reaction in which all NMPs were added simultaneously and incubated for 90 min; and a two-step reaction in which pyrimidines were added first for 60 min, followed by purines, with a 30 min incubation after each addition (**Figure 5A**). Notably, both reaction schemes can be performed in a single vessel (“one pot”). The PPK2-synthesized NTPs were then used to drive *in vitro* transcription of Pepper, an RNA aptamer that forms a fluorescence complex upon binding the substrate HBC530^45^.

**Figure 5.**
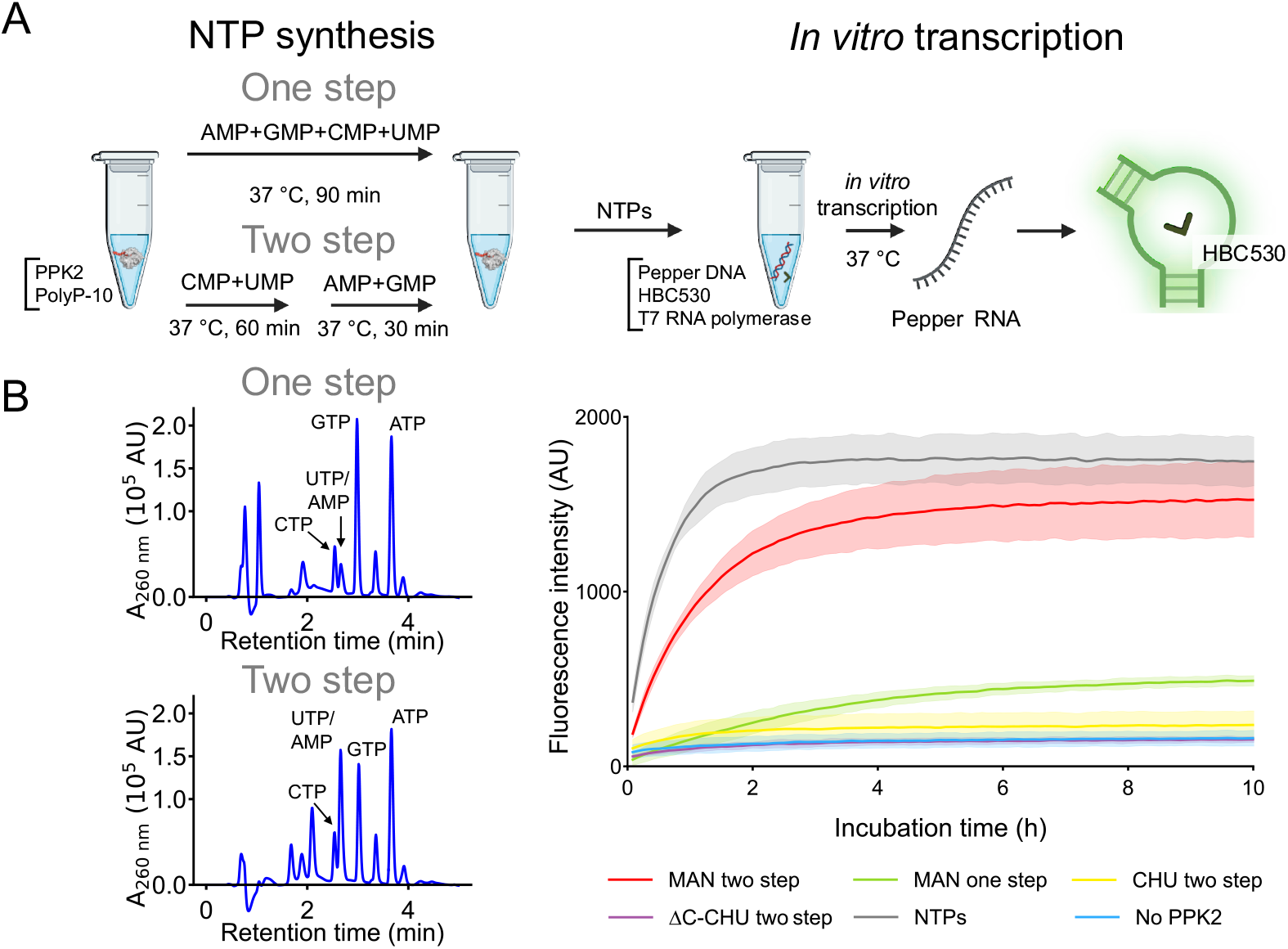
*In vitro* transcription utilizing NTPs produced by Class III PPK2 enzymes. **A**. NTP production and *in vitro* transcription reaction schemes. **B**. HPLC chromatograms for the one-step and two-step NTP production catalyzed by MAN (left). Time course of Pepper RNA production during *in vitro* transcription reactions with various Class III PPK2 enzymes (right). Plotted values are the average of 3 independent runs. Error bars are standard deviations.

The product distributions of the one-step and two-step reactions showed significant differences, particularly with respect to pyrimidines (**Figure 5B**). In the one-step reaction, CTP production could not be unambiguously assigned, and significant amounts of the UMP and CMP substrates remained. IVT reactions performed with this sample yielded modest transcription. The two-step reaction mixture, on the other hand, contained significant amounts of all four NTPs and the subsequent IVT reaction produced nearly as much RNA (90.5% yield), nearly as quickly, as the NTP positive control, with both reactions reaching completion at ∼4 h. The two-step reactions using CHU or truncated CHU (ΔC-CHU), on the other hand, failed to produce RNA even at the level of the one-step MAN reaction, highlighting the uniquely broad substrate specificity of the MAN enzyme.

## Conclusions

We demonstrate that MAN is a universal PPK capable of efficiently phosphorylating all eight common ribonucleotides – purines and pyrimidines, monophosphates and diphosphates – with PolyP. However, our analysis suggests that PPK2 enzymes in general (and MAN in particular) have a relatively rugged functional landscape, particularly for non-adenine bases. This conclusion follows from (a) the diverse properties of N138 Clade III PPK2 enzymes (**Figure 2**), (b) the idiosyncratic, non-uniform changes in substrate preference upon mutation (**Figures 4B** and **Figure S6**) and chimerization (**Figure 4C** and **E**), and (c) the general sensitivity of these enzymes to reaction conditions (**Figure 3**). Although the source of this complex behavior is unknown, several lines of evidence suggest that conformational dynamics may play a role. First, the C-terminal α-helix does not directly interact with the base, yet truncation of this α-helix consistently shifts substrate specificity to purines (**Figure S7**). Second, mutation of the PolyP binding site also alters substrate preferences (**Figure 4E**), again without directly interacting with the base of the nucleotide. Finally, the chimera constructs (**Figure 4C** and **Figure S9**) cannot be compactly described as transitions between MAN and CHU functional profiles, despite substituting all non-identical residues near the base-binding site. The coupling of conformational dynamics with function may also contribute to the abrupt transition from ATP formation to APn formation with increasing temperature.

Although the challenges ahead for detailed PPK2 functional prediction are significant, uPPK2 enzymes will be an exciting test case for investigating the structural determinants of broad substrate specificity. The biotechnological applications of uPPK2 enzymes are similarly exciting. We envision that MAN or enzymes like it – with the ability to phosphorylate all eight common ribonucleotides to NTP – will reduce the cost of DNA and RNA production and simplify the phosphorylation of non-canonical nucleotides. The one-pot nucleotide synthesis and *in vitro* transcription reaction scheme presented here marks a significant step towards the goal of harnessing PPK2 enzymes for human benefit.

## Materials and Methods

### Sequence analysis and gene tree construction

Full-length PPK2 gene sequences were retrieved from the InterPro database^46^ entry IPR022488 (Polyphosphate kinase-2-related) on 28 August 2022. Clustering and representative sequence selection were performed using CD-HIT version 4.8.1^47^ using a 60% identity cutoff and a word size of 4. The resulting 2,118 representative sequences were aligned using the L-INS-i algorithm implemented in MAFFT version 7.525^48^. To improve phylogenetic tree building, gaps and regions of poor alignment were removed using trimAL version 1.4^49^ using the gappyout option. Alignment quality was assessed visually in Jalview version 2.11.4^50^. A phylogenetic tree was constructed using IQ-TREE version 2.3.6 ^51^ with automatic model selection and a single thread, as multiple threads degrade performance^52^. Phylogenetic trees were analyzed using the python package Environment for Tree Exploration (ETE) version 4^53^ and visualized using the Interactive Tree of Life (iTOL) version 7.0 ^54^. PPK2 clades were assigned using the clade-specific residue at position 137, ^11,19^ and named according to established conventions.

### Cloning and mutagenesis

Class III PPK2 genes from *Arenitalea lutea* (ARE, A0A1M6FUJ9), *Leeuwenhoekiella palythoae* (LEE, A0A1M5Z1E8), *Mangrovibacterium marinum* (MAN, A0A2T5C445), and *Pedobacter sp*. (PED, A0A519×5M4), and their truncated C-terminal α-helix variants were synthesized (Eurofins Scientific) and cloned into the pQI-MBP plasmid. pQI-MBP, constructed in our group, is a derivative of the pQE30 plasmid (Qiagen) in which MBP is cloned downstream of the T5 promoter and also encodes the LacIq sequence. The full plasmid sequence is given in **Table S3**. The previously characterized PPK2 gene from *Cytophaga hutchinsonii* (CHU, A0A6N4SMB5)^19^ and its truncated C-terminal α-helix variant were also synthesized. Each construct contains an N-terminal 6xHis tag for purification followed by a maltose-binding protein (MBP) tag to improve expression and solubility. For the preparation of MAN mutants, site-directed mutagenesis was performed using the Q5 Site-Directed Mutagenesis Kit (New England Biolabs). Chimeras were prepared using the HiFi DNA Assembly Kit (New England Biolabs) according to the manufacturer’s instructions. All protein sequences are provided in **Table S2**.

### Expression and purification

PPK2 proteins were produced by heterologous expression in *E. coli* XL10-Gold (Agilent). Briefly, 1 L cultures of terrific broth (TB) were shake-incubated at 37 °C until exponential growth and an optical density at 600 nm between 0.4-0.8 AU. At this point, the temperature was decreased to 25 °C, and isopropyl-β-D-thiogalactopyranoside (IPTG) was added to a final concentration of 0.5 mM. Expression proceeded overnight for approximately 16 h. Cells were harvested by centrifugation at 6,000 × g for 10 min at 4 °C. Cell pellets were stored at -80 °C until lysis. Cells were resuspended in pre-chilled 50 mM HEPES-KOH pH 7.6, 300 mM NaCl, 0.01 mM DTT, and 10 % glycerol, re-pelleted as above and resuspended once again. Cells were lysed by sonication on ice, and the lysate was clarified by centrifugation at 20,000 × g for 30 min at 4 °C. Clarified lysate was incubated with TALON metal affinity resin (Takara Bio Inc.) for 30 min at 4 °C with gentle mixing. The resins were rinsed with 10 column volumes of 50 mM HEPES-KOH pH 7.6, 300 mM NaCl, 0.01 mM DTT, 15 % glycerol, and 10 mM imidazole and PPK2 was eluted from the beads in 50 mM HEPES-KOH pH 7.6, 300 mM NaCl, 0.01 mM DTT, 15 % glycerol, and 300mM imidazole Purified protein was passed through a PD-10 desalting column (Cytiva) pre-equilibrated with 50 mM HEPES-KOH pH 7.6, 100 mM KCl, 10 mM MgCl2, and 30% glycerol. Protein purity was assessed by SDS-PAGE using Coomassie Brilliant Blue stain. Protein concentrations were measured using A_280_ and calculated with extinction coefficients (ARE: 113220 M^-1^ cm^-1^, LEE: 111730 M^-1^ cm^-1^, MAN: 117230 M^-1^ cm^-1^, and PED: 121700 M^-1^ cm^-1^) estimated by Benchling (https://www.benchling.com/) based on the protein sequences. Purified protein samples were stored at -80 °C.

### Polyphosphate kinase activity assay

Polyphosphate kinase activity profiling against nucleoside monophosphates (NMPs) and nucleoside diphosphates (NDPs) was performed in 50 mM MOPS-NaOH pH 7.0, 10 mM MnCl2 at 37 °C. Reaction mixtures contained 65 mM *phosphate units*, 4 mM nucleotide, and 18.9 ^μ^M PPK2, and had a final volume of 20 ^μ^L. Three lengths of PolyP were tested: PolyP-10, PolyP-60, and PolyP-700 (Bioenex, Japan), which have an average of 12.5, 65, and 750 phosphate units per molecule, respectively. After 60 min, the samples were put on ice, the reactions were terminated by the addition of trichloroacetic acid (TCA) to a final concentration of 2.5%. The samples were then centrifuged at 20,000 × g for 30 min at 4 °C, which removed the protein component. The supernatant was analyzed by high-performance liquid chromatography (HPLC) as described below.

MAN temperature profiling was performed as above except that only AMP and PolyP-60 were tested, and the reaction temperature ranged from 20-55 °C. MAN metal profiling was performed as above except that only AMP and PolyP-60 were tested and 10 mM MgCl2, CaCl2, CoCl2, or NiCl2 were used in place of MnCl2. For chimeras, reaction mixtures contained 50 mM MOPS-NaOH pH 7.0, 10 mM MnCl_2_, 50 mM phosphate units of PolyP-60, 4 mM nucleotide, and 18.9 ^μ^M enzyme. For point mutants at position 138 (CHU numbering), reaction mixtures contained 50 mM Tris-HCl pH 8.0, 10 mM MgCl_2_, 20 mM phosphate units from PolyP-10, 4 mM nucleotide, and either 0.2 ^μ^M (AMP, ADP, GMP, and GDP) or 6 ^μ^M (UMP, UDP, CMP, and CDP) enzyme. Reactions were conducted at 30 °C for either 5 min (AMP, ADP, GMP, GDP, UMP, and UDP) or 10 min (CMP and CDP).

### High-performance liquid chromatography (HPLC)

The supernatants from TCA precipitation were diluted in two volumes of 50 mM triethylammonium acetate, 2 mM ethylenediaminetetraacetic acid, pH 7.0 (Buffer A). Samples were injected onto an XBridge BEH C18 column (2.5 ^μ^m particles, 4.6 × 30 mm, Waters Corporation) for analysis. Gradient elution was performed at 20 °C with a flow rate of 1 mL/min. Buffer A was the starting buffer and 100% acetonitrile served as the elution buffer (Buffer B). Buffer B was increased linearly from 0% to 2.5% over 2 min, and then further increased to 5% over the next 4 min. Absorbance at 260 nm was recorded. Nucleotide concentrations were determined from the peak areas of the chromatogram using LabSolutions version 5.124 (Shimadzu). Peak areas were converted into nucleotide concentrations according to a standard curve obtained with authentic nucleotides. Nucleotide extinction coefficients were obtained from Promega and NEB chemical data sheets.

### Analytical Ultracentrifugation (AUC)

Ultracentrifugation experiments were performed using a ProteoLab XLI analytical ultracentrifuge equipped with an AN-50 Ti rotor (Beckman Coulter). Protein samples with A_280_ ≅ 1 were prepared in 50 mM HEPES-KOH pH 7.6, 500 mM KCl, 10 mM MgCl2, and 0.005% Tween 20. 400 ^μ^L of each sample and reference buffer were injected into a 12 mm sample cell with a sapphire window. Velocity measurements were conducted at 50,000 rpm and 20 °C. For each sample, 100 scans were collected over the course of 6-12 h, depending on the number of samples being measured simultaneously. The density and viscosity of the buffer were calculated by SEDNTERP version 3.0.4^56^. Time course data was analyzed by SEDFIT version 16.50^57^ using a resolution of 100, a partial specific volume of 0.73, a frictional ratio of 1.2, and a confidence level of the F-ratio (f/fo) of 0.95.

### Steady-state kinetic analysis of MAN

Kinetic analysis of MAN was performed in 50 mM Tris-HCl buffer pH 8.0, 10 mM MgCl_2_, 20 mM phosphate units from PolyP-60 at 30 °C. To estimate initial reaction rates, the concentration of MAN and the sampling frequency were adjusted for each nucleotide and its concentration: AMP and GDP were varied from 1 to 20 mM, with MAN concentrations of 0.3 ^μ^M and 3 ^μ^M, respectively. For ADP, the substrate concentration was varied from 0.3 to 10 mM, with a MAN concentration of 1.2 ^μ^M. For GMP, the concentration ranged from 2.5 to 25 mM, with a MAN concentration of 1 ^μ^M. For UMP, UDP, and CMP, substrate concentrations were varied from 2.5 to 40 mM, with MAN concentrations of 1 ^μ^M, 3 ^μ^M, and 6 ^μ^M, respectively. The concentration of CDP was varied from 2.5 to 30 mM using a MAN concentration of 6 ^μ^M. Reaction times ranged from 6 min (AMP, ADP, GMP, and GDP) to 20 min (UMP, UDP, CMP, and CDP), and reactions were terminated by adding TCA to a final concentration of 2.5%. Initial rates were calculated from the linear regime of starting substrate depletion. For example, when the substrate is AMP, the initial rate was calculated from the AMP peak only, even if multiple products (*i*.*e*., ADP, ATP, or APn) are present. Finally, *k*_cat_ and K_M_ were obtained by fitting initial reaction velocity data to the Michaelis-Menten equation. All analyses were performed in Python using SciPy version 1.13.1.

#### Class III PPK2-powered *in vitro* transcription

*In vitro* transcription (IVT) was monitored by fluorescence of the Pepper RNA aptamer (**Table S4**) upon binding to benzylidene-cyanophenyl (HBC) derivatives. The *in vitro* reaction mixture was composed of 40 mM Tris-HCl buffer pH 8.0, 50 mM NaCl, 8 mM MgCl_2_, 5 mM dithiothreitol, 0.25 mM spermidine, 0.001% bovine serum albumin (BSA), 62 ng of Pepper DNA, 0.04 ^μ^M pyrophosphatase (derived from yeast ^58^and purified in-house), 10 ^μ^M HBC530 (which binds to the Pepper RNA aptamer), 0.8 units of T7 RNA polymerase from the ScriptMAX Thermo T7 Transcription Kit (Toyobo), and nucleotides prepared as described below. The positive control contained 0.125 mM of each NTP, whereas test samples contained 0.125 mM nucleotide mixtures derived from a one-pot reaction with PPK2. The PPK2 reaction mixture contained 18.9 ^μ^M PPK2 enzyme in 50 mM Tris-HCl, 10 mM MgCl_2_, 30 mM PolyP-10, and 1 mM NMPs (AMP, GMP, CMP, and UMP). In the one-step reaction, all NMPs were added and incubated for 90 min at 37 °C. In the two-step reaction, CMP and UMP were added and the sample was incubated for 60 min. Then AMP and GMP were added, and the sample was incubated for an additional 30 min. The PPK2 reaction mixtures were used as a source of nucleotides for the IVT. IVT was performed at 37 °C for 20 h in an Mx3005P real-time PCR thermocycler (Agilent). Pepper RNA production was tracked by measuring fluorescence with excitation and emission wavelengths of 492 nm and 516 nm, respectively.

## Acknowledgments

This study was supported by the Human Frontier Science Program Grant Number RGP003/2023 (TM), JSPS KAKENHI Grant Numbers 22K21344 (TM), and 21H05228 (TM).

