## Supplementart Information for "A universal polyphosphate kinase powers *in vitro* transcription"

[1]Department of Life Science and Technology, Institute of Science Tokyo, Tokyo, Japan
[2]Earth-Life Science Institute, Institute of Science Tokyo, Tokyo, Japan
[3]Blue Marble Space Institute of Science, Seattle, USA
[4]The authors contributed equally to this study
†Current address: MAQsys Inc., Kanagawa, Japan

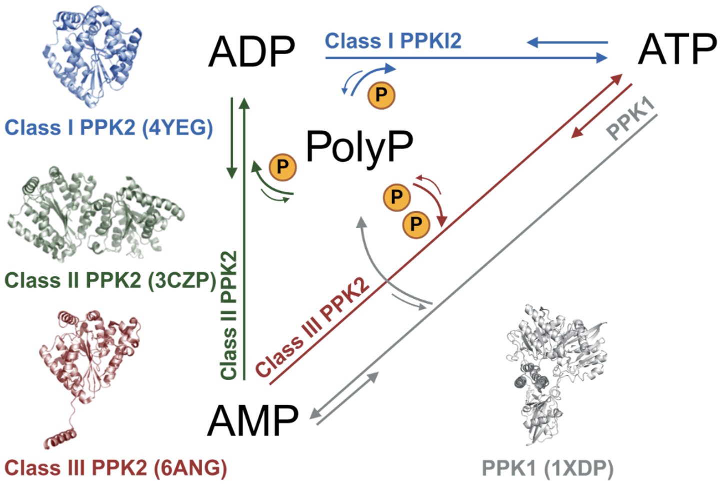


**Figure S1**. Schematic showing PPK catalytic diversity. Both PPK1 and PPK2 use PolyP and nucleotides as substrates. PPK2 is divided into three classes in accordance with the phylogenetic tree. Catalytic profiles may be not strictly distinguished by the class.

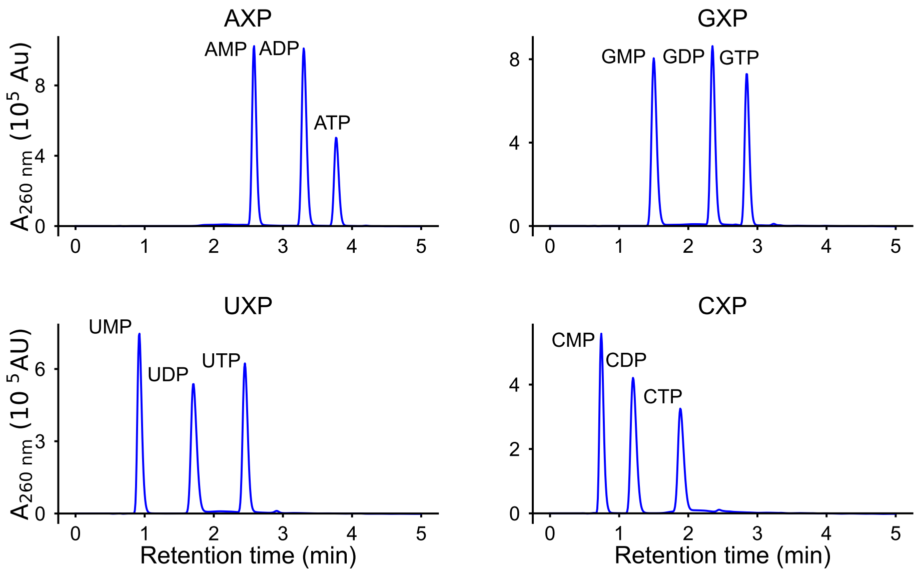


**Figure S2**. HPLC analysis of nucleotides. "X" is inclusive of mono- (M), di- (D), and tri- (T) phosphate forms. For each base, the nucleotides with different degrees of phosphorylation elute as distinct peaks with almost no overlap. All nucleotides were quantified under the same conditions by HPLC. Here, the results for authentic samples of 333 μM of each nucleotide (999 μM total) are shown.

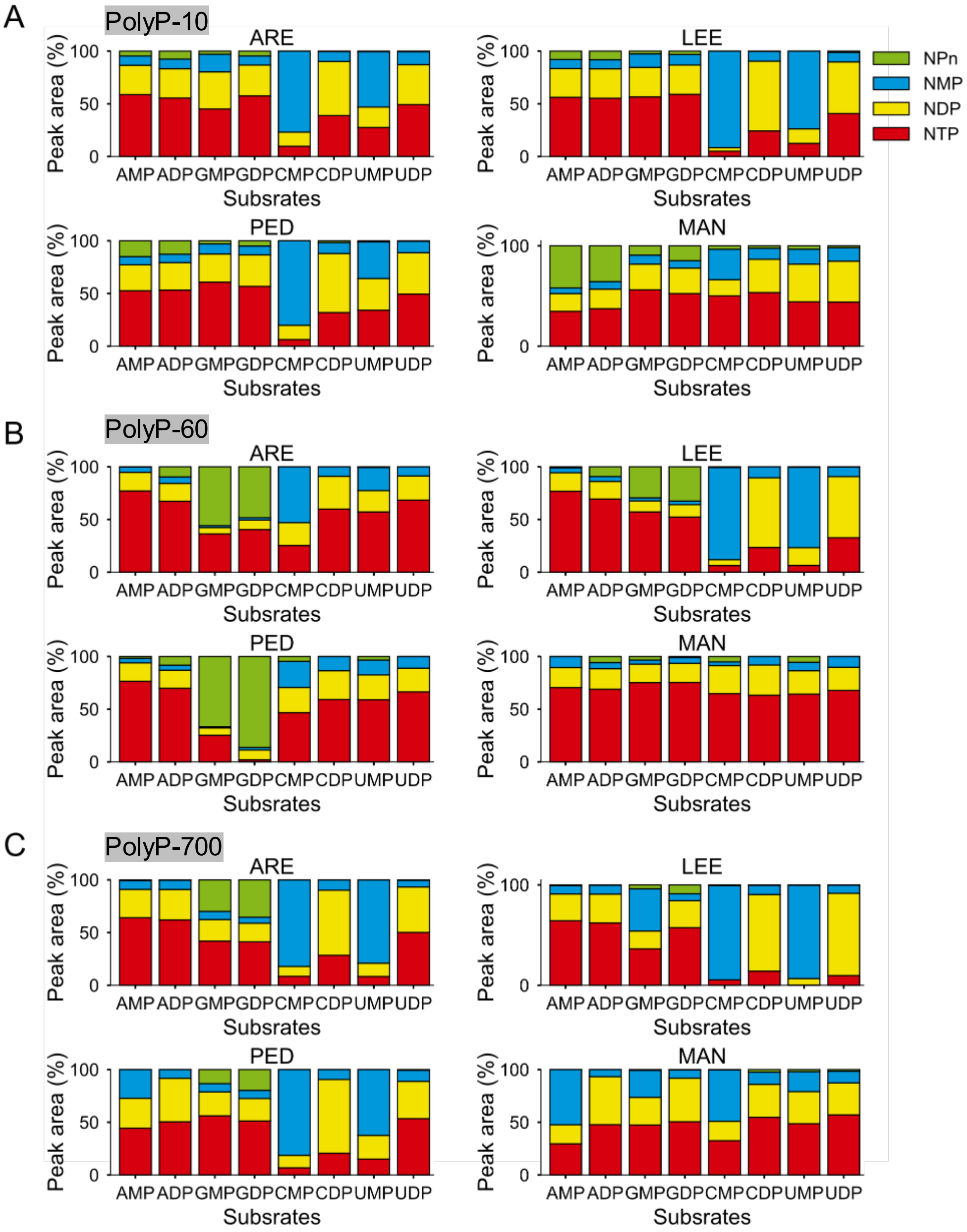


**Figure S3**. NMP and NDP phosphorylation activity of Class III PPK enzymes (ARE, LEE, PED, and MAN) using PolyP with different lengths. **Figure 2B** reports these same data as NTP synthesis efficiency, while the bar plots here show the product distribution with (A) Poly10, (B) PolyP50, and (C) PolyP700.

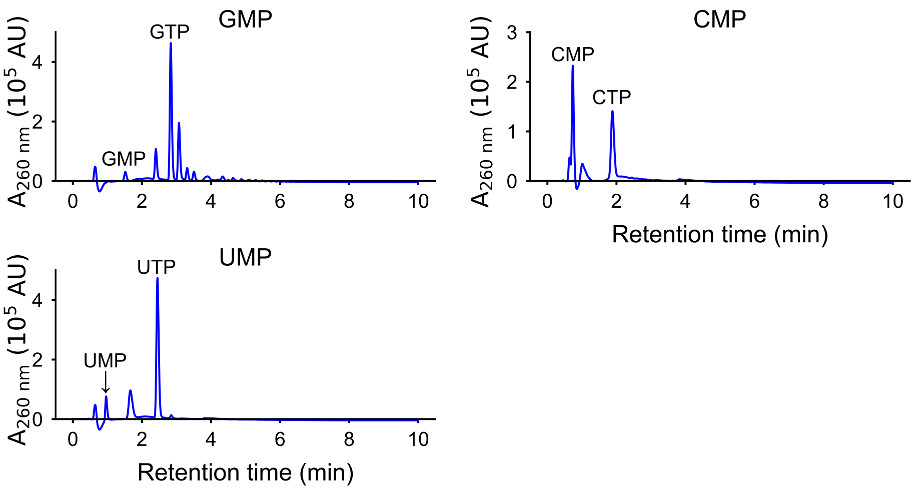


**Figure S4**. Polyphosphate nucleotides generated by MAN. The reaction was carried out at 55 °C as in **Figure 3A** except that GMP, CMP, or UMP was used as donor the substrate instead of AMP. While some minor over-phosphorylation of GMP was observed, neither of the pyrimidine nucleotides were significantly over-phosphorylated.

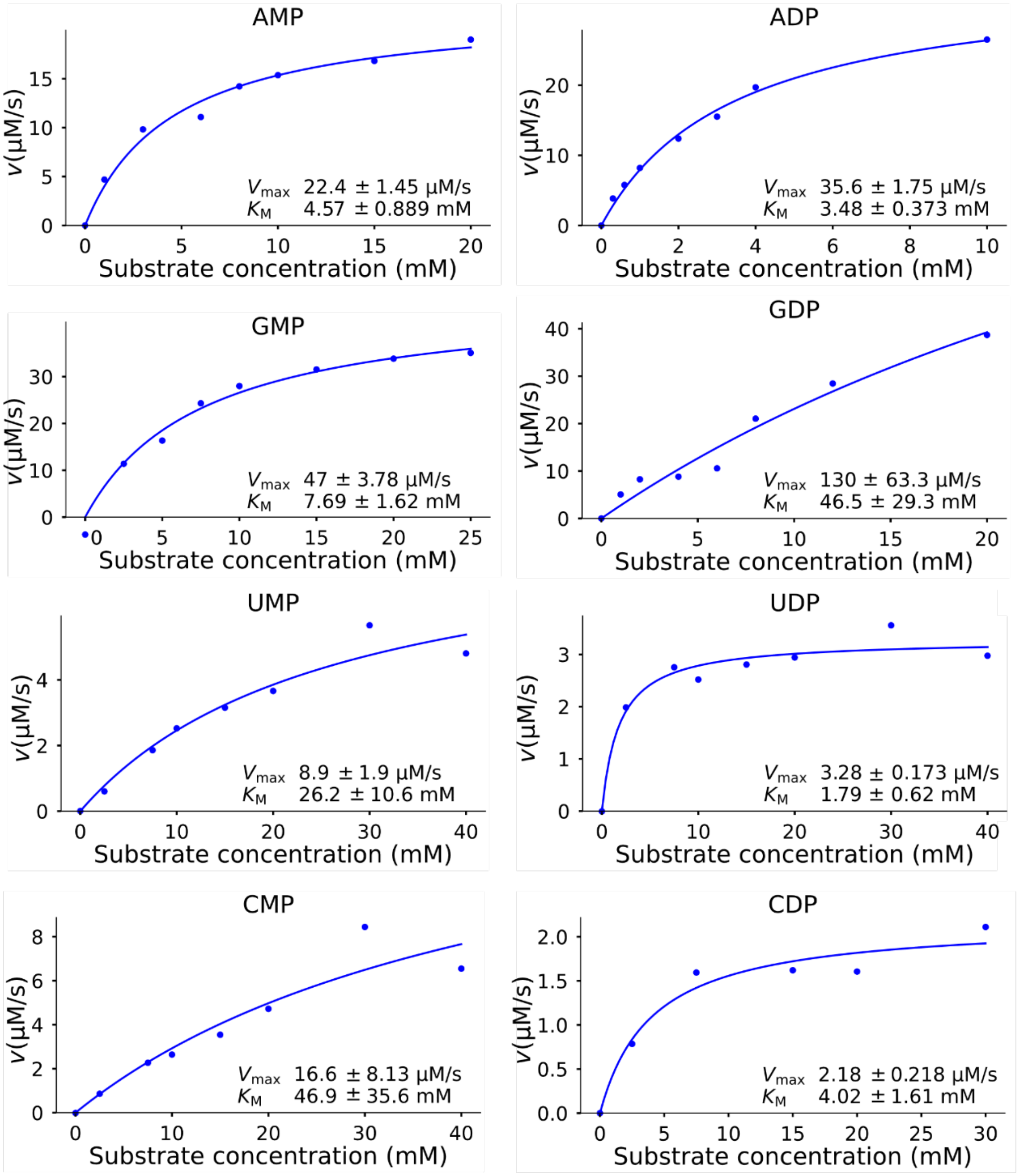


**Figure S5**. Michaelis-Menten plots from which $k_{cat}$ and $K_M$ values were determined. See also **Table 1**.

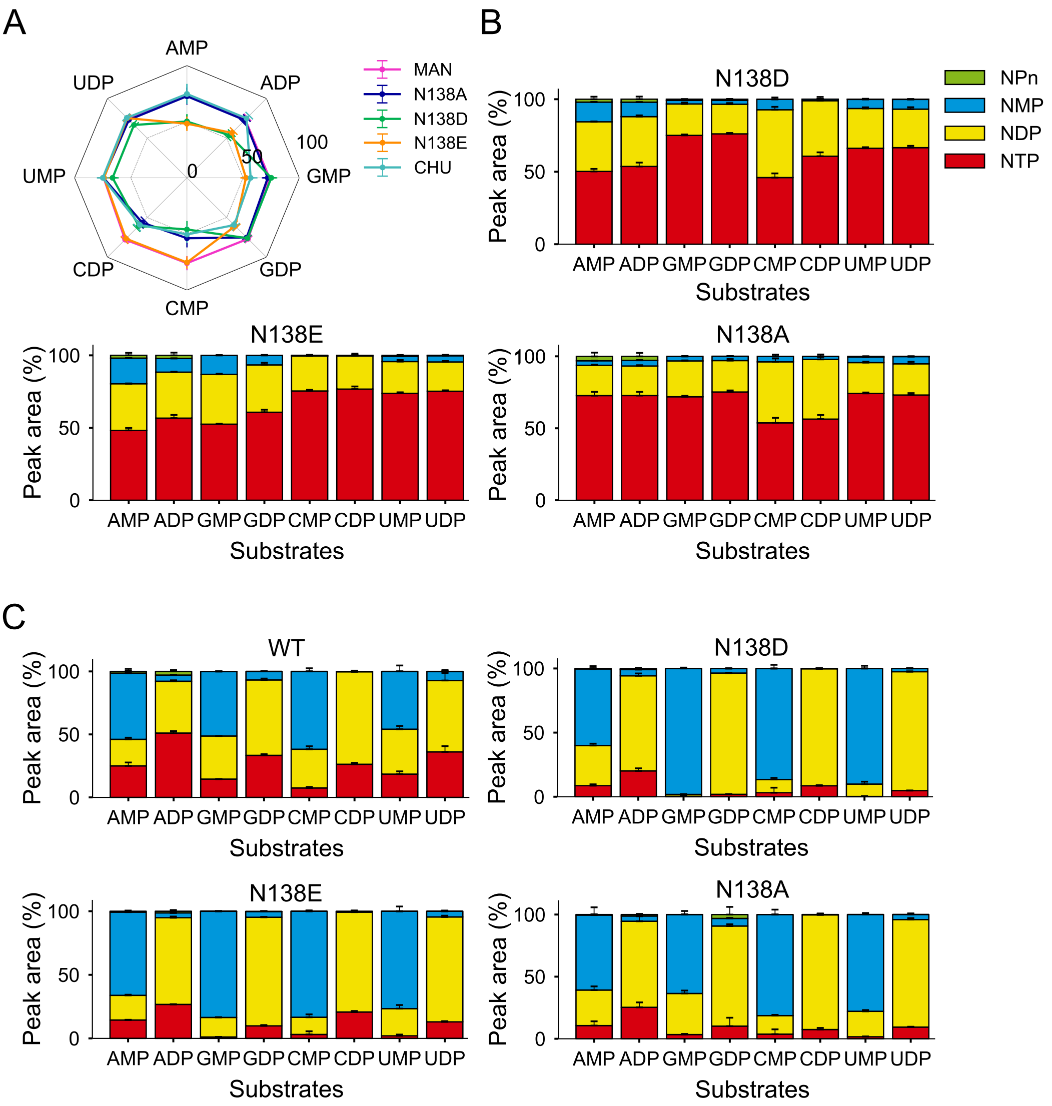


**Figure S6**. N138 point mutant endpoint activity assay. See also **Figure 4A**. (A) Percent conversion to NTP for enzymes (10 µM) reacting with the indicated nucleotide (4 mM) in the presence of 10 mM $Mn^{2+}$. (B) Product distributions for the high concentration mutant endpoint activity assay are shown in panel A. (C) Product distributions for the high concentration mutant. See **Figure 4B** for a plot of NTP synthesis efficiency. Plotted values are the average of 4 independent runs. Error bars are standard deviations.

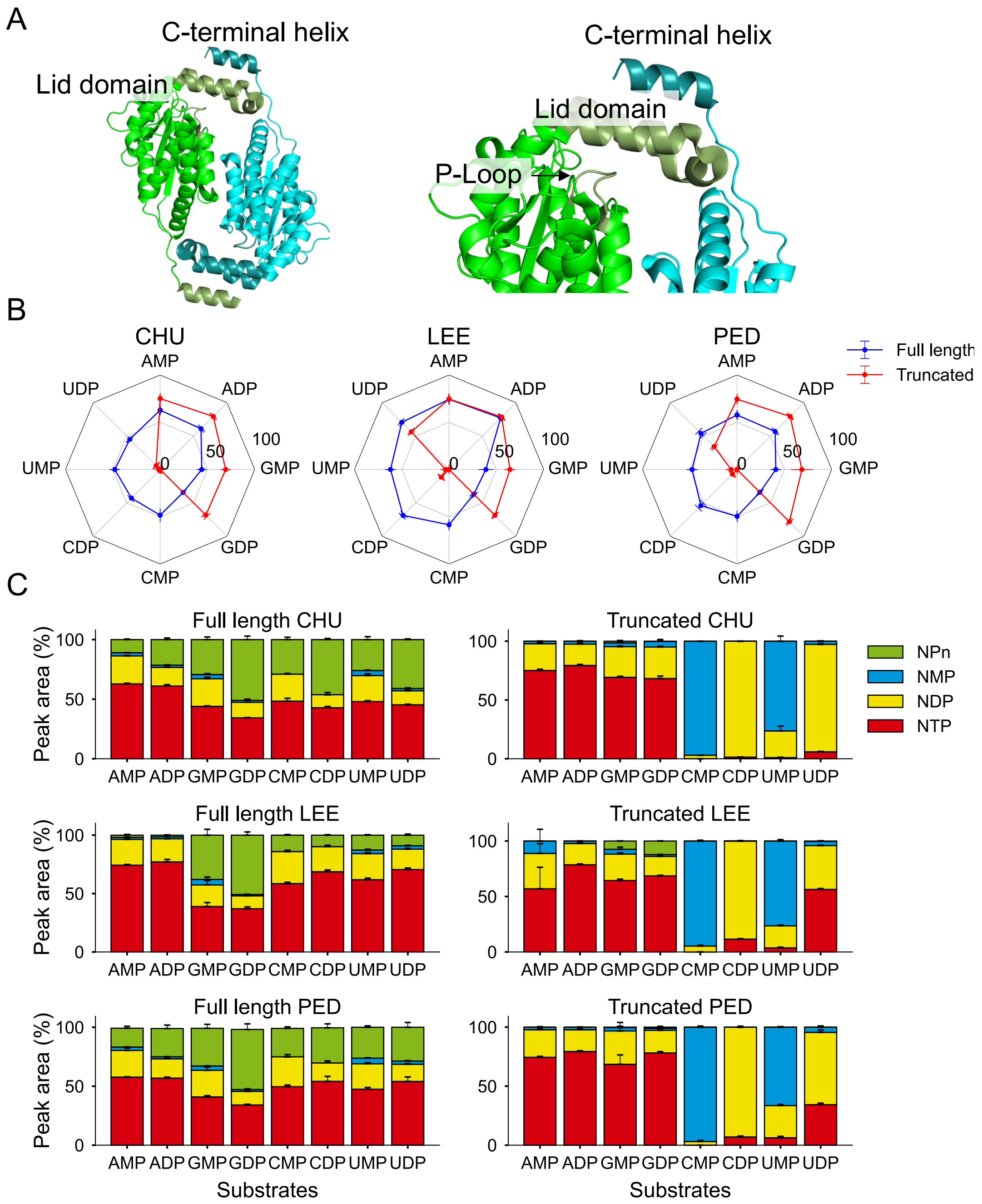


**Figure S7**. Influence of C-terminal α-helix on the activity of Class III PPK2. (A) The C-terminal helix of one protomer rests one layer away from the active site P-loop of another protomer (left) (PDB ID 6aqe). As such, the C-terminal helix may influence the P-loop conformations or dynamics through the lid domain (right) (PDB ID 6aqe). Thus, truncating the sequence may affect the nucleotide phosphorylation. Previously, it was shown that the phosphorylation of GMP and GDP was improved by the C-terminal α-helix truncation of CHU [1]. (B) Percent conversion to NTP for enzymes reacting with the indicated nucleotide (4 mM) in the presence of 10 mM $Mn^{2+}$ using full-length (blue) and C-terminal α-helix truncated (red) CHU, LEE, and PED. The reaction was performed with 18.9 μM enzyme, 20 mM PolyP-10, and 4 mM of indicated substrate in the standard reaction buffer. (C) The product distributions of reactions are analyzed in (B). Plotted values are the average of 4 independent runs. Error bars are standard deviations.

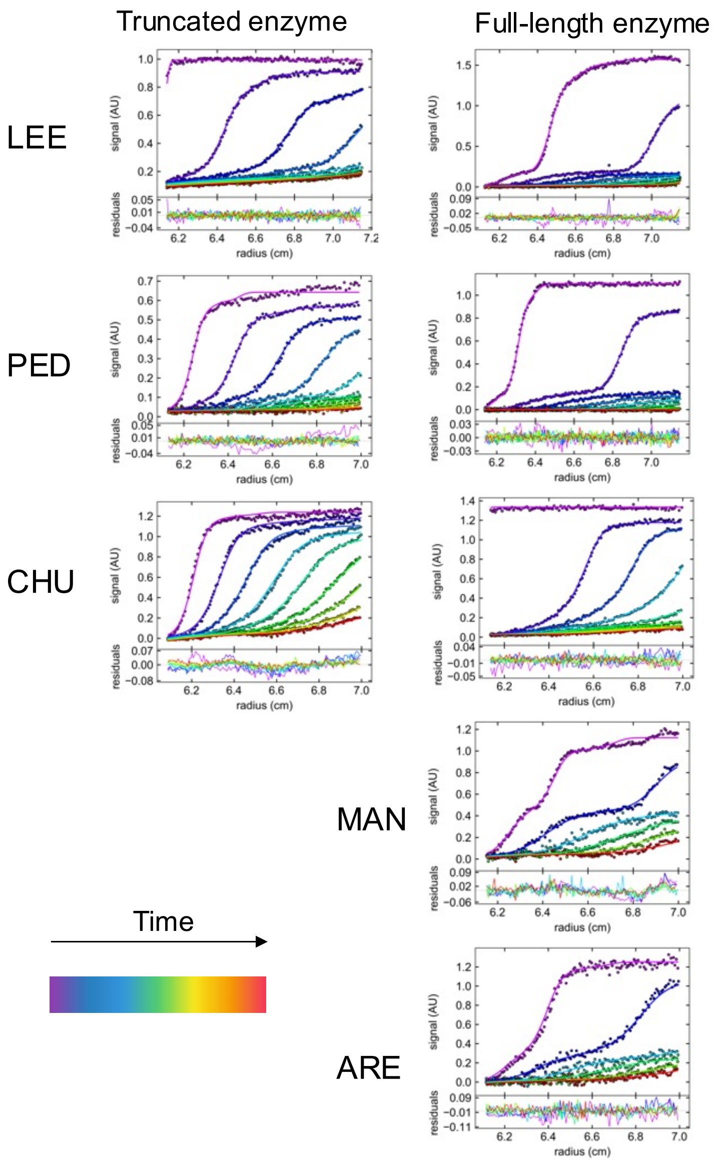


**Figure S8**. Analytical ultracentrifugation (AUC) data for full-length LEE, PED, CHU, MAN, and ARE (right) and C-terminal α-helix truncated LEE, PED, and CHU (left). Data of truncated MAN and ARE were not obtained due degradation in *E. coli* and low purification yields. Raw data (scatterplot) and fitted curves (lines) of absorbance scans at 280 nm as a function of radius for each time point are presented. Calculated molecular weights are given in **Table S1**. Briefly, the truncation of the C-terminal α-helix altered the oligomerization state of the PPK2 enzymes: While the full-length enzymes were a dimer, truncation of the C-terminus converted the enzymes into a monomer.

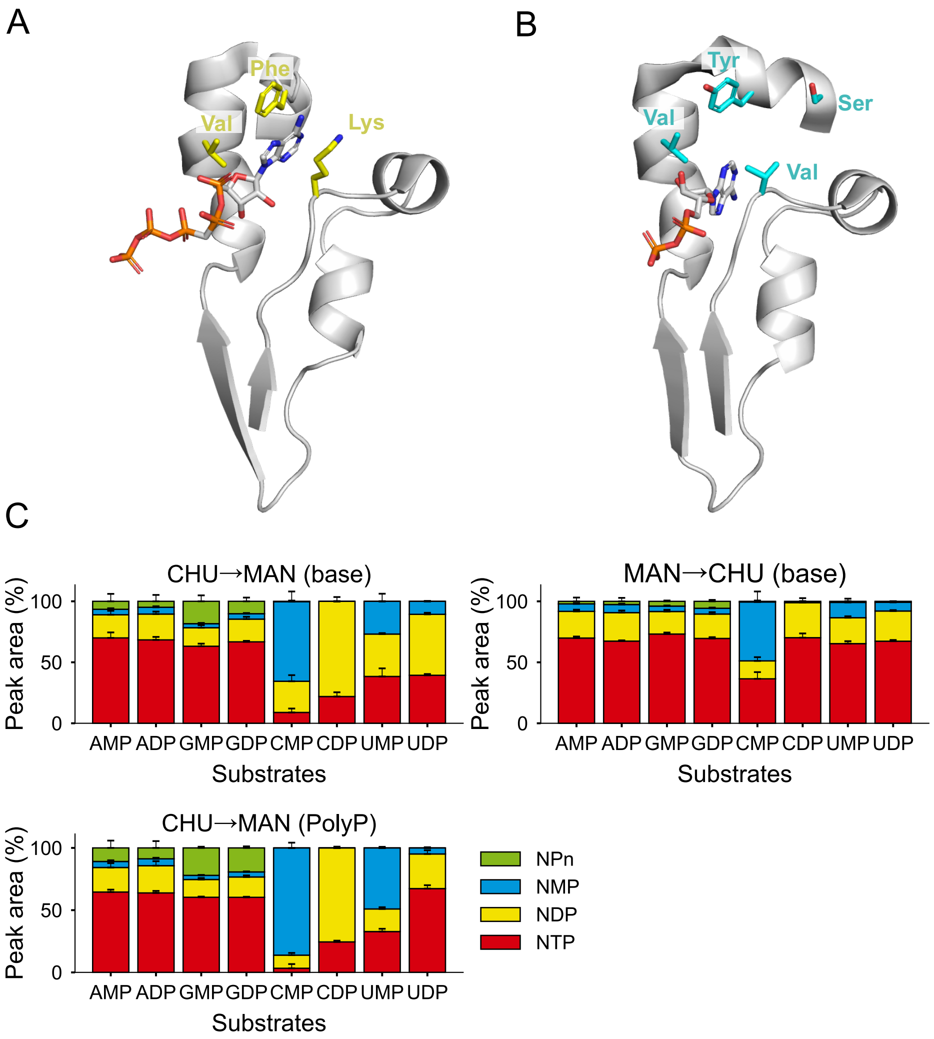


**Figure S9.** NMP and NDP phosphorylation activity of MAN mutants. Nucleotide binding modes for the Class I PPK2 from *Francisella tularensis* (FRA) bound to adenosine-5′-[(β,γ)-methyleno]pentaphosphate [2] (5LLB, panel A) and for CHU bound to ADP (6AN9, panel B) [1]. MAN-CHU chimerized binding site positions (**Figure 4**) are shown in sticks and colored. Note that the serine of CHU has no equivalent position in FRA. The base binding position of FRA is flipped up relative to CHU, allowing for direct interaction with the chimerized binding site positions than occurs in the canonical binding mode. (C) Product distributions of nucleotide phosphorylation by MAN or CHU chimeras. See **Figure 4C** and 4**E** for NTP synthesis efficiency. Plotted values are the average of 4 independent runs. Error bars are standard deviations.

**Table S1**. Molecular weight and inferred oligomerization state of PPK2 enzymes by analytical ultracentrifugation. Raw data is presented in **Figure S9**.

| Enzyme | Molecular weight by AUC (kDa) | Molecular weight theoretical (kDa) | $MW_{AUC}$ / $MW_{theoretical}$ |
|---|---|---|---|
| Truncated LEE | 88.8 ± 0.2 | 74.5 | 1.2 |
| Truncated PED | 89.0 ± 0.2 | 75.1 | 1.2 |
| Truncated CHU | 84.9 ± 0.5 | 75.9 | 1.1 |
| Full-length LEE | 179.6 ± 0.2 | 76.5 | 2.3 |
| Full-length MAN | 190.0 ± 0.5 | 78.0 | 2.4 |
| Full-length PED | 179.7 ± 0.2 | 76.8 | 2.3 |
| Full-length CHU | 159.3 ± 0.4 | 78.3 | 2.0 |
| Full-length ARE | 180.7 ± 0.7 | 76.3 | 2.4 |

**Table S2**. Enzyme amino acid sequences.

| Name | Uniprot ID | Protein sequence[#,##] |
|---|---|---|
| MAN | A0A2T5<br>C445 | MKSKQLNTDLFKVQAKQQIRLKNYDPASCNGFANRKEAEHQLKQDIKELA<br>DLQYQLYAENKRSVLIVFQAMDAAGKDGSIRHVFSGLNPQGCKVYSFKSP<br>SANELDHDYLWRHYNKLPPRGEIGIFNRSHYENVLITRVHPEFLLNENLP<br>DVHATNDITTEFWERRYRQIRDFEQTLTENGTTIIKFFLHLSKDEQKKRF<br>MERIRKPAKNWKFSSADIRERQYWDDYQHAYEQAMRATSTEEAPWYIIPA<br>DNKWFTHVAIGNIIVETLQQMNIQMPEISSEEKDALKKAKKKLQNE |
| PED | A0A519<br>X5M4 | MKTNTDKYLVAAKTKVKLKNFDTDYNGHLDKEAGKAELESVKEELSKFQE<br>VLYASNAHSLLIIFQAMDAAGKDGAIAHVMSGLNPQGCQVYSFKTPSSEE<br>YDHDFLWRHYKALPEKGRIGIHNRSHYENVLVCKVHPEYVLTENIPGYDD<br>VKKIDDNFWKGRYESIRNFEKHLAANGVAILKFFLHVSKDEQKQRFLDRI<br>EDPAKNWKFSAGDISERALWDDYMSAYEDALSETSTEDAPWYVIPADKKW<br>YTRLAVSQIIAEKLKNLNLEFPKLPKEETDALEGYKKQLLSEK |
| LEE | A0A1M<br>5Z1E8 | MKASDYQITAPIKLADLKTHEDFGKTKELEQELDEVSKKIAAIQNAMYAH<br>GKYAALICIQGMDTAGKDSLIREVFKNFNARGVVVHSFKTPTSLELGHDY<br>LWRHYIALPARGKVGVFNRTHYENVLVTRVHPEYILNENIPGVNDVSDID<br>EAFWEKRFEQINNFEKHIADNGTIIFKFFLHLSKEEQKHRLLRRLNKPNK<br>NWKFSAGDLKERALWDEYQQCYEDAINKTSKFHAPWYIIPADDKPTARLI<br>LANILLEKLSSYKDIEEPELDEKTKSNIDAYRKQLESE |
| ARE | A0A1M<br>6FUJ9 | MNDYKVTSNISLKNTQTKVVIDDAEKQLKKLRKELGKLQDTLYAHGKYSV<br>LVCLQGMDTAGKDSLIREVFKDFNARGVVVHSFKVPTELELKHDYLWRHY<br>IALPARGKFGVFNRTHYENVLVTRVHPYYIMGENIPGVESVEDVNDAFWN<br>KRFEQINNFEKHIAENGTIIFKFFLNLSKAEQKNRLLRRLRKQEKNWKFS<br>PGDLKERKLWDKYQNCYEDAINRTSKPHAPWFTIPADDKPTARYLVAKIM<br>LDTLKTYNNIKEPELDDDIKANIEMYKQQLNNE |

[#]Underlined residues correspond to the C-terminal α-helix.

[##]N-terminal MBP tag not shown. For full constructs, see **Table S3**.

**Table S3**. Full plasmid sequence for MAN.

| Sequence of plasmid pQI-MBP# |
|---|
| 5′tggcgaatgggacgcgccctgtagcggcgcattaagcgcggcgggtgtggtggttacgcgcag |
| cgtgaccgctacacttgccagcgccctagcgcccgctcctttcgctttcttcccttcctttctcg |
| ccacgttcgccggctttccccgtcaagctctaaatcgggggctccctttagggttccgatttagt |
| gctttacggcacctcgaccccaaaaaacttgattagggtgatggttcacgtagtgggccatcgcc |
| ctgatagacggtttttcgccctttgacgttggagtccacgttctttaatagtggactcttgttcc |
| aaactggaacaacactcaaccctatctcggtctattcttttgatttataagggattttgccgatt |
| tcggcctattggttaaaaaatgagctgatttaacaaaaatttaacgcgaattttaacaaaatatt |
| aacgtttacaatttcaggtggcacttttcggggaaatgtgcgcggaacccctatttgtttatttt |
| tctaaatacattcaaatatgtatccgctcatgagacaataaccctgataaatgcttcaataatat |
| tgaaaaaggaagagtatgagtattcaacatttccgtgtcgcccttattcccttttttgcggcatt |
| ttgccttcctgtttttgctcacccagaaacgctggtgaaagtaaaagatgctgaagatcagttgg |
| gtgcacgagtgggttacatcgaactggatctcaacagcggtaagatccttgagagttttcgcccc |
| gaagaacgttttccaatgatgagcacttttaaagttctgctatgtggcgcggtattatcccgtat |
| tgacgccgggcaagagcaactcggtcgccgcatacactattctcagaatgacttggttgagtact |
| caccagtcacagaaaagcatcttacggatggcatgacagtaagagaattatgcagtgctgccata |
| accatgagtgataacactgcggccaacttacttctgacaacgatcggaggaccgaaggagctaac |
| cgcttttttgcacaacatgggggatcatgtaactcgccttgatcgttgggaaccggagctgaatg |
| aagccataccaaacgacgagcgtgacaccacgatgcctgcagcaatggcaacaacgttgcgcaaa |
| ctattaactggcgaactacttactctagcttcccggcaacaattaatagactggatggaggcgga |
| taaagttgcaggaccacttctgcgctcggcccttccggctggctggtttattgctgataaatctg |
| gagccggtgagcgtgggtctcgcggtatcattgcagcactggggccagatggtaagccctcccgt |
| atcgtagttatctacacgacggggagtcaggcaactatggatgaacgaaatagacagatcgctga |
| gataggtgcctcactgattaagcattggtaactgtcagaccaagtttactcatatatactttaga |
| ttgatttaaaacttcatttttaatttaaaaggatctaggtgaagatcctttttgataatctcatg |
| accaaaatcccttaacgtgagttttcgttccactgagcgtcagaccccgtagaaaagatcaaagg |
| atcttcttgagatcctttttttctgcgcgtaatctgctgcttgcaaacaaaaaaaccaccgctac |
| cagcggtggtttgtttgccggatcaagagctaccaactctttttccgaaggtaactggcttcagc |
| agagcgcagataccaaatactgtccttctagtgtagccgtagttaggccaccacttcaagaactc |
| tgtagcaccgcctacatacctcgctctgctaatcctgttaccagtggctgctgccagtggcgata |
| agtcgtgtcttaccgggttggactcaagacgatagttaccggataaggcgcagcggtcgggctga |
| acggggggttcgtgcacacagcccagcttggagcgaacgacctacaccgaactgagatacctaca |
| gcgtgagctatgagaaagcgccacgcttcccgaagggagaaaggcggacaggtatccggtaagcg |
| gcagggtcggaacaggagagcgcacgagggagcttccagggggaaacgcctggtatctttatagt |
| cctgtcgggtttcgccacctctgacttgagcgtcgatttttgtgatgctcgtcaggggggcggag |
| cctatggaaaaacgccagcaacgcggcctttttacggttcctggccttttgctggccttttgctc |
| acatgttctttcctgcgttatcccctgattctgtggataaccgtattaccgcctttgagtgagct |
| gataccgctcgccgcagccgaacgaccgagcgcagcgagtcagtgagcgaggaagcggaagagcg |
| cctgatgcggtattttctccttacgcatctgtgcggtatttcacaccgcatatatggtgcactct |
| cagtacaatctgctctgatgccgcatagttaagccagtatacactccgctatcgctacgtgactg |
| ggtcatggctgcgccccgacacccgccaacacccgctgacgcgccctgacgggcttgtctgctcc |
| cggcatccgcttacagacaagctgtgaccgtctccgggagctgcatgtgtcagaggttttcaccg |
| tcatcaccgaaacgcgcgaggcagctgcggtaaagctcatcagcgtggtcgtgaagcgattcaca |
| gatgtctgcctgttcatccgcgtccagctcgttgagtttctccagaagcgttaatgtctggcttc |
| tgataaagcgggccatgttaagggcggttttttcctgtttggtcactgatgcctccgtgtaaggg |
| ggatttctgttcatgggggtaatgataccgatgaaacgagagaggatgctcacgatacgggttac |
| tgatgatgaacatgcccggttactggaacgttgtgagggtaaacaactggcggtatggatgcggc |
| gggaccagagaaaaatcactcagggtcaatgccagcgcttcgttaatacagatgtaggtgttcca |
| cagggtagccagcagcatcctgcgatgcagatccggaacataatggtgcagggcgctgacttccg |
| cgtttccagactttacgaaacacggaaaccgaagaccattcatgttgttgctcaggtcgcagacg |
| ttttgcagcagcagtcgcttcacgttcgctcgcgtatcggtgattcattctgctaaccagtaagg |
| caaccccgccagcctagccgggtcctcaacgacaggagcacgatcatgcgcacccgtggggccgc |
| catgccggcgataatggcctgcttctcgccgaaacgtttggtggcgggaccagtgacgaaggctt |

```
gagcgagggcgtgcaagattccgaataccgcaagcgacaggccgatcatcgtcgcgctccagcga
aagcggtcctcgccgaaaatgacccagagcgctgccggcacctgtcctacgagttgcatgataaa
gaagacagtcataagtgcggcgacgatagtcatgccccgcgcccaccggaaggagctgactgggt
tgaaggctctcaagggcatcggtcgagatcccggtgcctaatgagtgagctaacttacattaatt
gcgttgcgctcactgcccgctttccagtcgggaaacctgtcgtgccagctgcattaatgaatcgg
ccaacgcgcggggagaggcggtttgcgtattgggcgccagggtggtttttcttttcaccagtgag
acgggcaacagctgattgcccttcaccgcctggccctgagagagttgcagcaagcggtccacgct
ggtttgccccagcaggcgaaaatcctgtttgatggtggttaacggcgggatataacatgagctgt
cttcggtatcgtcgtatcccactaccgagatatccgcaccaacgcgcagcccggactcggtaatg
gcgcgcattgcgcccagcgccatctgatcgttggcaaccagcatcgcagtgggaacgatgccctc
attcagcatttgcatggtttgttgaaaaccggacatggcactccagtcgccttcccgttccgcta
tcggctgaatttgattgcgagtgagatatttatgccagccagccagacgcagacgcgccgagaca
gaacttaatgggcccgctaacagcgcgatttgctggtgacccaatgcgaccagatgctccacgcc
cagtcgcgtaccgtcttcatgggagaaaataatactgttgatgggtgtctggtcagagacatcaa
gaaataacgccggaacattagtgcaggcagcttccacagcaatggcatcctggtcatccagcgga
tagttaatgatcagcccactgacgcgttgcgcgagaagattgtgcaccgccgctttacaggcttc
gacgccgcttcgttctaccatcgacaccaccacgctggcacccagttgatcggcgcgagatttaa
tcgccgcgacaatttgcgacggcgcgtgcagggccagactggaggtggcaacgccaatcagcaac
gactgtttgcccgccagttgttgtgccacgcggttgggaatgtaattcagctccgccatcgccgc
ttccactttttcccgcgttttcgcagaaacgtggctggcctggttcaccacgcgggaaacggtct
gataagagacaccggcatactctgcgacatcgtataacgttactggtttcacattcaccaccctg
aattgactctcttccgggcgctatcatgccataccgcgaaaggttttgcgccattcgatggtgtc
cgggatctcgacgctctcccttatgcgactcctgcattaggaagcagcccagtagtaggttgagg
ccgttgagcaccgccgccgcaaggaatggtgcatgcaaggagatggcgcccaacagtcccccggc
cacggggcctgccaccatacccacgccgaaacaagcgctcatgagcccgaagtggcgagcccgat
cttccccatcggtgatgtcggcgatataggcgccagcaaccgcacctgtggcgccggtgatgccg
gccacgatgcgtccggcgtagaggatcgagatctcgatcccgcgaaattaatacgactcactata
ggggaattgtgagcggataacaattcccctctagaaataattttgtttaactttaagaaggagat
atacatatggctagcatgactggtggacagcaaatgggtcgcggatccgaattcgaggccctgag
ggccATGAAATCTAAGCAACTTAATACTGACTTGTTCAAGGTTCAAGCCAAGCAGCAGATCCGCT
TGAAGAACTATGATCCCGCCTCCTGTAACGGTTTTGCCAACCGCAAAGAGGCGGAACATCAATTA
AAGCAGGACATCAAAGAACTTGCAGACCTGCAGTACCAACTTTACGCAGAGAATAAACGTAGTGT
CTTAATTGTGTTTCAAGCCATGGATGCCGCTGGAAAAGACGGTAGTATCCGTCATGTGTTTTCGG
GCTTAAACCCACAGGGTTGTAAGGTGTACTCCTTCAAAAGTCCGAGTGCGAACGAATTGGACCAC
GATTATTTATGGCGCCATTACAACAAGTTGCCGCCACGCGGCGAGATTGGTATCTTTAACCGCAG
CCACTACGAGAATGTTCTGATTACACGTGTACATCCTGAATTCTTACTGAACGAGAATTTGCCCG
ATGTTCACGCCACAAATGACATTACTACGGAATTTTGGGAGCGTCGTTACCGTCAAATCCGCGAC
TTTGAACAGACCCTTACAGAAAATGGTACTACAATTATTAAATTCTTTTTACATTTGTCTAAGGA
CGAACAAAAAAAACGTTTCATGGAACGCATCCGTAAACCGGCAAAAAATTGGAAGTTCTCCTCTG
CTGACATCCGCGAACGTCAGTATTGGGATGATTATCAACATGCCTACGAGCAGGCTATGCGCGCG
ACCAGTACCGAAGAAGCCCCATGGTACATTATCCCCGCCGATAACAAGTGGTTCACGCACGTTGC
GATTGGCAACATCATCGTAGAAACTCTTCAACAGATGAACATTCAGATGCCGGAGATCTCATCGG
AAGAAAAGGATGCTCTGAAAAAAGCTAAAAAGAAGTTACAAAATGAGgccgcactcgagcaccac
caccaccaccactgagatccggctgctaacaaagcccgaaaggaagctgagttggctgctgccac
cgctgagcaataactagcataaccccttggggcctctaaacgggtcttgaggggttttttgctga
aaggaggaactatatccggat
```

[#] MAN sequence is indicated with capital letters.

**Table S4**

| |
|---|
| Sequence of RNA10-Pepper fusion construct[#,##] |
| 5′GGGTCCCGGCGCCAGTGCCTGCCGAAGCAGGCCGACACGCCACGATTGGGGACCCtttTGGTC<br>ATGTGATCGGCGTATGCACACGGCACGCGAATAGCACAACACCCTGCGATCTCGCCGGGCAGTTA<br>AACAGCGAGTTCGGCCGACAAAGTATCGTGTCATCGTCGTCCCTATAGTGAGTCGTATTA |

[#]T7 promoter sequence underlined.

[##]The sequence preceding 'ttt' corresponds to the Pepper sequence and the sequence following 'ttt' corresponds to the RNA10 and T7 RNA promoter sequences. RNA10 is reported to bind a phospholipid membrane[3]. RNA10-Pepper fusion sequence was used for IVT.
